# Timing Control of Purkinje Cell Outputs by Dual cAMP Actions on Axonal Action Potential and Transmitter Release

**DOI:** 10.1101/2023.09.14.557678

**Authors:** Kei Furukawa, Takuma Inoshita, Shin-ya Kawaguchi

## Abstract

All-or-none digital signaling based on high-fidelity action potentials (APs) in neuronal axons is pivotal for the temporally precise sending of identical outputs rapidly to widespread multiple target cells. However, technical limitation to directly measure the signaling in small size of intact axonal structures has hindered the evaluation of high-fidelity signal propagation. Here, using direct recordings from axonal trunks and/or terminals of cerebellar Purkinje cells in culture and slice, we demonstrate that the timing of axonal output is delayed by the second messenger cAMP without clear changes of transmission efficacy. Slowed axonal signaling upon cAMP increase was ascribed to negative control of axonal Na^+^ channels, leading to smaller and hence slower conduction of APs specifically at an axon. On the other hand, a facilitatory effect of cAMP on presynaptic transmitter release, which generally operates at various CNS synapses, was also evident as augmented release probability in Purkinje cell axon terminals, compensating for weakening of release by the reduction of Ca^2+^ influx upon smaller AP. Taken these results together, our *tour-de-force* functional dissection of inhibitory axonal signaling unveiled a dynamic control of synaptic output timing by cAMP keeping output strength constant.

## Introduction

Neuronal computation relies on axonal digital signaling with regenerative action potentials (APs) which enable rapid information transfer over long distance (Llinás, 1988). Classically APs are regarded as all-or-none type of rigid and faithful signals in terms of strength and velocity, and hence in principle unsusceptible for modification. Such characteristics of APs are defined by the axonal membrane excitability reflected by enrichment of Na^+^ channels relative to K^+^ channels together with the passive electrical property of membrane determined by cytoplasmic axial resistance and membrane resistance (Coombs et al., 1955). The concept of high-speed and precise signaling in the nervous system based on the digital APs lies at the center of understanding how neurons work. However, recent studies are changing the classical view of such a rigorous axonal AP conduction (Goaillard et al., 2020).

This conceptual advance was critically due to the finding of an analogue signaling capability of axons, that is, AP sometimes changes its amplitude, time-course, and/or even conduction velocity (Debanne et al., 2011; Shu et al., 2006; Zbili and Debanne, 2019). However, evaluation of axons’ physiology is demanding because of technical hurdle for direct electrophysiological recordings from a distal compartment of an intact axon (as small as ∼ μm) hundreds μm away from the soma. Exceptionally, recent elegant subcellular patch-clamp recording of APs from a tiny axonal compartment demonstrated speeding-up or -down of AP conduction in response to an increase in intracellular second messenger, cAMP, in cerebellar excitatory neurons (Byczkowicz et al., 2019) or in the cortical pyramidal cells (Lezmy et al., 2021), respectively. Such changes of AP conduction velocity are ascribed to the altered membrane excitability at axon, which could be accompanied by changes of AP waveforms, the resultant Ca^2+^ influx, and transmitter release from terminals. Thus, the AP conduction velocity and the waveform, which defines Ca^2+^ influx causing transmitter release, are likely in nature tightly related. Nevertheless, it remains elusive whether the two essential factors of APs, conduction velocity and waveforms, can be independently modulated or not. In addition, cAMP is also known to positively modulate presynaptic transmitter release (Byrne and Kandel, 1996; Capogna et al., 1995; Chavez-Noriega and Stevens, 1994; Chen and Regehr, 1997; Huang et al., 1994; Kaneko and Takahashi, 2004; Meadows et al., 2021; Saitow et al., 2000; Salin et al., 1996; Trudeau et al., 1996; Weisskopf et al., 1994), suggesting a possibility that AP conduction change by cAMP might be accompanied with changes in synaptic output strength through modulation of presynaptic release machinery.

To study the axonal rapid propagation of APs at a high temporal resolution together with biophysical analysis of presynaptic transmitter release machinery, direct patch clamp recordings from intact long axons/terminals are essential. The axon and terminal of cerebellar Purkinje cells (PCs) could be an ideal neuronal model, which allows to perform such a *tour-de-force* functional dissection (Kawaguchi and Sakaba, 2015). Taking advantage of that technical merit, we quantitatively measured how the AP propagation and the resultant transmitter release is modulated by cAMP increase in a PC axon, and found that AP conduction velocity is lowered through shaping the AP waveform, which is accompanied by decreased membrane excitability via reduction of axonal Na^+^ current. We also examined how the attenuated AP and cAMP pathway cooperatively affect the Ca^2+^-triggered transmitter release form a PC bouton.

## Results

### Delayed synaptic outputs from a PC axon by cAMP

To evaluate how cAMP affects axonal signaling in a GABAergic cerebellar PCs, we first examined the effects of cAMP on synaptic outputs from a PC in culture, in which long axonal arborization (sometimes beyond 1 mm) is preserved from the soma to terminals on target cells, unlike acute slice preparation. To specifically visualize PCs, we used an adeno-associated virus (AAV) vector, AAV2-CA-EGFP that preferentially infects PCs among neurons in the cerebellar culture, as in previous studies (Kawaguchi and Sakaba, 2015). Simultaneous whole-cell somatic patch-clamp recordings were performed from a presynaptic PC and its postsynaptic target neuron, in the presence of the AMPA receptor antagonist NBQX (10 μM) (Figure 1A). Presynaptic PCs were current-clamped so that the membrane potentials were around -70 mV, and APs were elicited by current injection to the PC soma (500 - 900 pA for 10 ms), which evoked IPSCs (eIPSCs) in the voltage-clamped (at – 80 to -100 mV) postsynaptic target cell (Figure 1B). eIPSCs showed large fluctuation in amplitude (in average 896 ± 317 pA), consistent with previous studies (Kawaguchi and Sakaba, 2015). To increase cytoplasmic cAMP, we added the cell-permeable adenylyl cyclase activator forskolin (50 μM) to the external bath. In clear contrast to previous studies showing cAMP-induced facilitation of transmission at excitatory synapses (Chavez-Noriega and Stevens, 1994; Chen and Regehr, 1997; Huang et al., 1994; Salin et al., 1996; Weisskopf et al., 1994), unexpectedly, the eIPSC amplitude was not significantly different from the basal condition after forskolin application (90% ± 9% at 3 min; Figure 1B, C). On the other hand, we observed a remarkable increase in the synaptic delay after the forskolin application, estimated from a latency between the AP peak to the onset of the eIPSC, corresponding to the sum of two temporal parameters for AP conduction time and the subsequent exocytosis of synaptic vesicles (114 ± 2%, p < 0.001; 1.96 ± 0.22 ms for 518 ± 85 μm conduction to 2.28 ± 0.27 ms; Figure 1E, F). Forskolin altered neither the 20-80% risetime of eIPSCs (0.54 ± 0.03 ms to 0.56 ± 0.03 ms, p = 0.562) nor the coefficient of variation of their amplitudes (CV, 0.39 ± 0.06 to 0.45 ± 0.07, p = 0.449) (Figure 1D). Furthermore, the somatic AP waveforms were little affected by forskolin except for a tendency of slight attenuation in size accompanied with marginally slower kinetics probably through lowering membrane resistance (see Supplementary Figure 1). These results suggest that cAMP increases the latency of synaptic output in a PC without affecting its strength.

**Figure 1.**
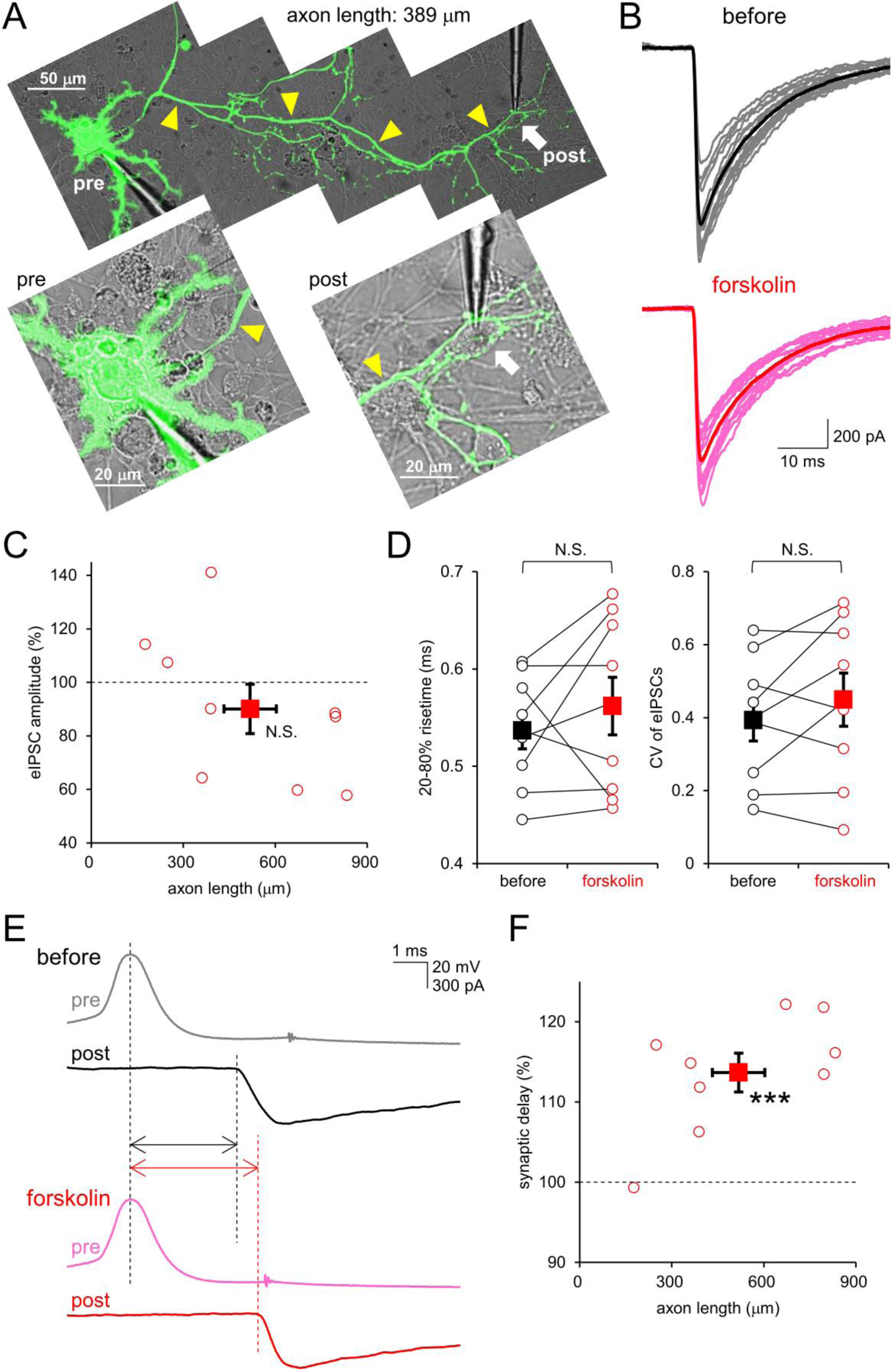
**A,** Images of an EGFP-labeled cultured PC. Paired whole-cell recorded presynaptic PC soma (lower left) and its postsynaptic target neuron (upper right, white arrow) are shown in magnified images. The length of PC’s axon (yellow arrowheads) from the presynaptic soma to the postsynaptic cell was 389 μm. **B**, Representative traces of eIPSCs (gray, pink) and their average (black, red) before (top) and after (bottom) forskolin application (50 μM). **C**, **D**, Normalized averaged amplitudes (C), 20-80% risetime (D, left) and CV (D, right) of eIPSCs for individual cells (open circles) and mean ± SEM (closed squares) after (connected by lines) forskolin application. n = 9, p = 0.108. Risetime, p = 0.354; CV, p = 0.203. **E**, Representative traces of APs (gray, pink) and the corresponding eIPSCs (black, red) before (top) and after (bottom) application of forskolin. Traces are aligned by the peak time of presynaptic APs. Synaptic delay (double arrows) is highlighted. **F**, Normalized synaptic delays averaged in individual cells (open circles) and mean ± SEM (closed squares) after forskolin application plotted against the axonal length between two cells. n = 9, p = 0.0001

### Slowing down of AP conduction by cAMP in a PC axon

Recent studies demonstrated opposite directions of cAMP-mediated modulation of the axonal AP conduction velocity: speeding up in the cerebellar cortical circuit such as mossy and parallel fibers (Byczkowicz et al., 2019) and slowing down in cortical pyramidal cells (Lezmy et al., 2021). We hypothesized that cAMP might modulate the AP conduction also along a PC axon in a negative direction, and performed simultaneous paired recordings of spontaneous APs from both a PC soma and its axon (Figure 2A and Supplementary Figure 2A). EGFP fluorescence enabled us to precisely target a thin patch pipette to the axonal trunk. By measuring delays between the spontaneous AP peaks at the soma and axon, we evaluated the AP conduction velocity (Figure 2B). Supporting our idea, paired cell-attached recordings demonstrated that forskolin increased the time needed for the AP to reach the axonal recording site (465 ± 49 μm away) (150 ± 9%, p < 0.01), indicating that cAMP slows down the AP conduction in a PC axon (540 ± 80 μm/ms to 380 ± 50 μm/ms; Figure 2B, C). The increase in time for AP conduction tended to be larger as the distance from the soma to the recorded axonal site, although not significant (Figure 2C). In addition, the axonal cell-attached AP waveform, but not that at the soma, tended to be slower in time course and smaller in amplitude after the forskolin application (somatic cell-attached AP amplitude: 99 ± 5%, p = 0.71; axonal cell-attached AP amplitude: 58 ± 4%, p < 0.001; Figure 2B and Supplementary Figure 2), implying that the AP conduction in axon, but not that around the soma, might be negatively regulated by the increase in cytoplasmic cAMP. The above demonstrated action of cAMP on a PC axon is similar to that in a cortical pyramidal cell (Lezmy et al., 2021), and quite contrasts to the previous findings of increased AP conduction velocity by cAMP on mossy and parallel fibers in the cerebellum (Byczkowicz et al., 2019).

**Figure 2.**
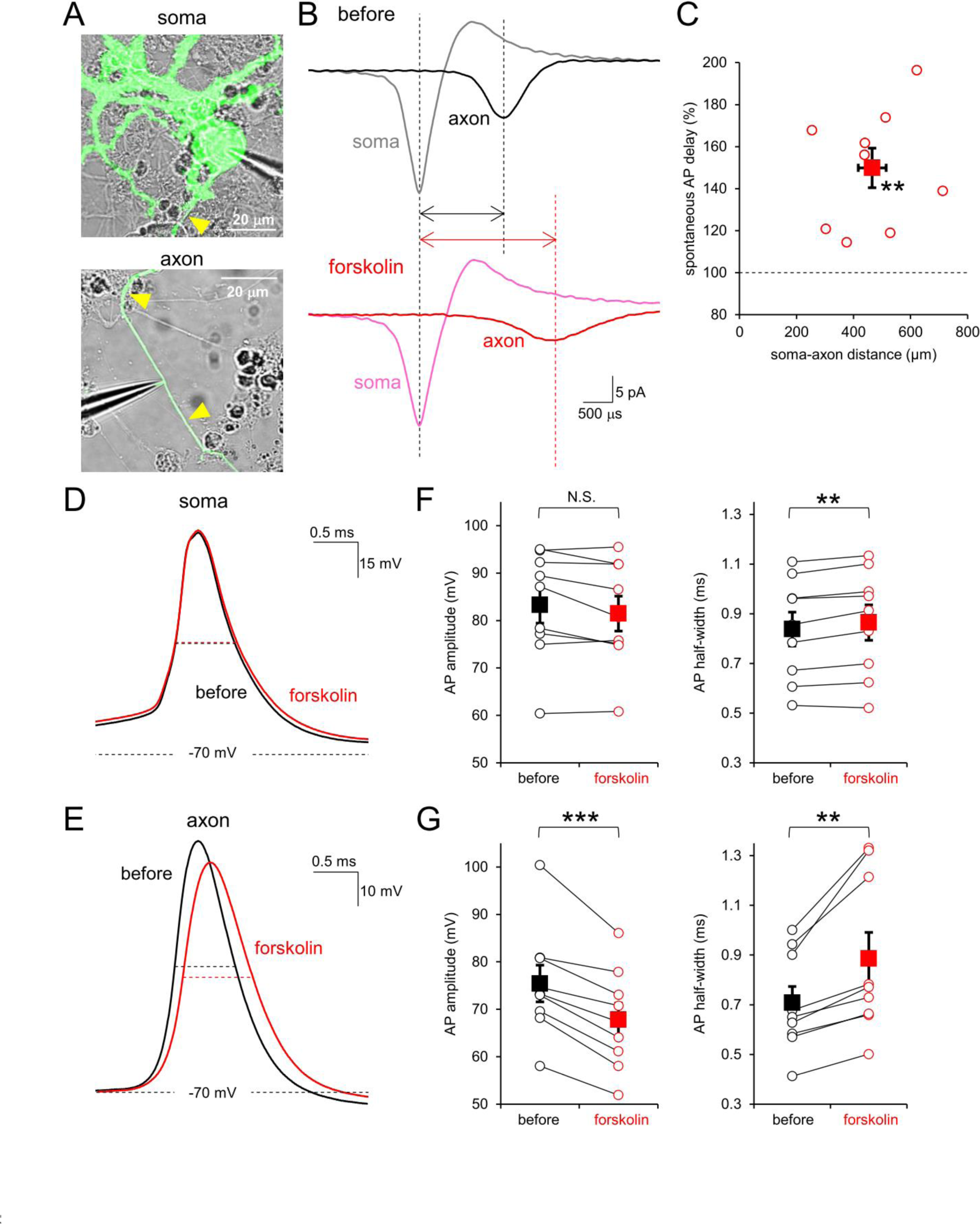
**A,** Images of pair-recorded soma (top) and axon (bottom) of an identical PC. Yellow arrowheads show the axon. **B**, Representative averaged traces of cell-attached recorded AP at soma (gray, pink) and axon (black, red) before (top) and after (bottom) forskolin application. APs traces are aligned by the peak of somatic APs. The AP delay is highlighted by double arrows. **C**, Normalized AP delays averaged in individual cells (open circles) and mean ± SEM (closed squares) after forskolin application plotted against the soma to axonal distances. n = 9, p = 0.0023. **D**, **E**, Representative averaged AP traces at the soma (D) and axon (E) elicited by somatic current injection before (black) and after (red) forskolin application. Half-widths are shown as dotted lines. **F**, **G**, Average amplitudes (left) and half-widths (right) of APs at soma (F) and axon (G) of individual cells (open circles), and mean ± SEM (closed squares) before and after forskolin application. n = 9 cells. Soma: amplitude, p = 0.055; half-width, p = 0.0039; Axon: amplitude, p = 0.00016; half-width, p = 0.0031

### Attenuation of APs by cAMP-mediated excitability change in a PC axon

To understand the mechanism by which the AP conduction velocity is negatively regulated by cAMP in a PC axon, we focused on the somatic and axonal AP waveforms during propagation in terms of their susceptibility to cAMP modulation. The soma and axon were whole-cell current-clamped so that the membrane potentials were around -70 mV except for the time triggering AP firings by current injection to the soma (Figure 2D). As in non-invasive cell-attached recordings (shown in Figure 2B, C), the slowing down of AP conduction by forskolin was evident in paired current-clamp recordings (Supplementary Figure 3A). Importantly, an axon exhibited remarkable reduction in amplitude and slowed time course of AP waveforms after the forskolin application (amplitude: 90 ± 1%, p < 0.001; halfwidth: 122 ± 3%, p < 0.001; Figure 2E, G), in contrast to the little effects on the somatic AP (except for a marginal, but significant slowing of time course) (amplitude: 99 ± 1%, p = 0.059; halfwidth: 103 ± 1%, p < 0.01; Figure 2D, F), suggesting that cAMP attenuates AP waveforms during conduction. Taken together, our direct patch clamp recordings from an axon demonstrated negative regulation of APs by cAMP in size and propagation velocity selectively in an axon.

To explore the molecular mechanism for the axonal AP modulation by cAMP, next we directly recorded voltage-activated currents from a PC axon (Figure 3). Under the voltage-clamped configuration, the axonal membrane was depolarized for tens of ms by step pulses from -70 mV, and inward and/or outward currents through voltage-gated channels were recorded. As shown in Figure 3A, B (and Supplementary Figure 4), depolarization at axonal membrane caused a large transient inward current (-2.13 ± 0.30 nA peaking at -40 mV) corresponding to Na^+^ influx, and the following sustained outward currents through K^+^ channels (4.25 ± 0.43 nA at 20 mV), in consistence with previous studies (Kawaguchi and Sakaba, 2015). Surprisingly, application of forskolin significantly reduced the amplitudes of Na^+^ currents, but not K^+^ currents, without changing the voltage sensitivity (Na^+^ currents: -1.72 ± 0.20 nA at -40 mV; K^+^ currents: 3.63 ± 0.41 nA at 20 mV, respectively; Figure 3A, B and Supplementary Figure 4A and 5). Similarly, cytoplasmic dialysis of the cyclic AMP analogue, cAMPS-Sp administrated from a patch pipette also decreased Na^+^ currents, but not K^+^ currents (Na^+^ currents: -1.76 ± 0.27 nA peaking in control and -0.99 ± 0.16 nA with cAMPS-Sp, p = 0.033; K^+^ currents: 4.23 ± 0.46 nA at 20 mV in control and 3.42 ± 0.82 nA with cAMPS-Sp, p = 0.409; Supplementary Figure 4C and Supplementary Table 1). These results indicate that the membrane excitability, mainly determined by the relative strength of Na^+^ influx against K^+^ efflux, is clearly decreased by the cytoplasmic cAMP increase in a PC axon. Interestingly, cAMP elevation by forskolin had no effect on the somatically-recorded I_Na_^+^ (-4.51 ± 0.76 nA to -4.32 ± 0.74 nA at -20 mV; -2.31 ± 1.05 nA to -2.17 ± 0.92 nA at 0 mV, p = 0.29 and 0.56, respectively). In addition, we also recorded the hyperpolarization-activated currents (I_h_), cation influx typically through HCN channels, by applying hyperpolarization pulses for 300 ms. In quite contrast to the previous studies suggesting cAMP-mediated changes of I_h_ currents in the cerebellar parallel and mossy fibers (Byczkowicz et al., 2019), forskolin little affected the amplitude of I_h_ in a PC axon (Figure 3C, D), probably due to the selective expression of cAMP-insensitive HCN1 in PCs (Notomi and Shigemoto, 2004; Wang et al., 2001). Thus, among voltage-dependent channels tested here, Na_v_ channels were specifically and negatively regulated by the cytoplasmic cAMP increase in a PC axon, leading to lower membrane excitability, which would negatively control the AP conduction characterized with attenuated amplitude, slower time course, and lower conduction velocity.

**Figure 3.**
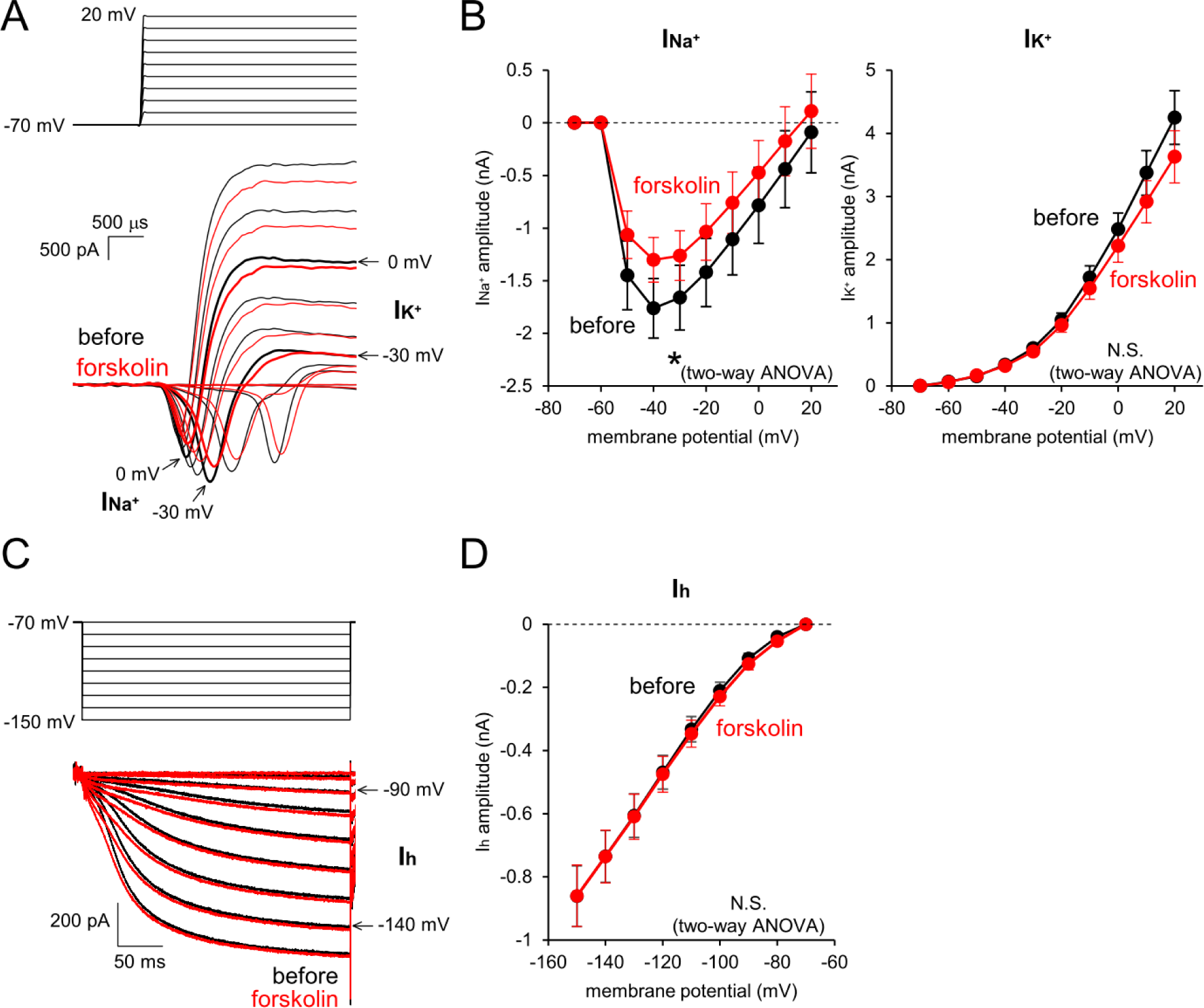
**A,** Representative voltage-gated Na^+^ and K^+^ currents recorded from a PC axon (bottom) upon depolarization steps from -70 mV to +20 mV (top). **B**, Current-voltage relations for axonal Na^+^ (left) and K^+^ (right) currents before (black) and after (red) forskolin application. Data are presented as mean ± SEM. n = 7 axons. Na^+^ currents, p = 0.029; K^+^ currents, p = 0.071. **C**, Representative voltage-gated currents recorded from a PC axon (bottom) upon hyperpolarization steps from -70 mV to -150 mV (top). **D**, Current-voltage relations for HCN currents before (black) and after (red) forskolin application. Data are presented as mean ± SEM. n = 7 axons, p = 0.749.

Our results so far indicate that cAMP attenuates the AP conduction mainly through suppressing Na^+^ channels, leading to slow timing of synaptic outputs from the terminals. What is the cAMP’s downstream molecule involved? To address this, we applied various pharmacological agents and tested whether the reduction of Na^+^ current by cAMP elevation is affected or not. We performed a direct recording from a PC axon, and the voltage-dependent Na^+^ currents were measured in the presence of H89 (PKA inhibitor, 1 μM) or ESI-09 (Epac inhibitor, 100 μM). Even in the presence of H89 or ESI-09, the Na^+^ influx was reduced upon the forskolin application (H89: -2.01 ± 0.31 nA to -1.37 ± 0.31 nA, p < 0.001; ESI-09: -2.07 ± 0.36 nA to -1.65 ± 0.43 nA, p < 0.05; Supplementary Figure 4D, E), suggesting that the negative regulation of Na^+^ channels by cAMP elevation are dependent neither on PKA nor Epac.

### Augmentation of release by cAMP in a PC terminal

Our above results suggested that cAMP attenuates AP amplitudes and slows down the conduction velocity probably through negative modulation of Na^+^ channels. Attenuation of AP at a PC axon terminal was previously shown to reduce the Ca^2+^ influx and the subsequent GABA release (Diaz-Rojas et al., 2015; Kawaguchi and Sakaba, 2015). Thus, the lack of clear change in amplitudes of eIPSCs after forskolin application (shown in Figure 1) might imply some compensatory mechanism to augment release, counteracting the reduced AP amplitudes at the terminals. Indeed, many types of neurons show augmentation of presynaptic transmitter release after the application of forskolin (Midorikawa and Sakaba, 2017; Yao and Sakaba, 2010), particularly at excitatory synapses with relatively low release probability based on the loose coupling of release machinery to presynaptic Ca^2+^ channels.

To evaluate the effect of cAMP on release machinery in a PC terminal, we first monitored spontaneous miniature IPSCs (mIPSCs) from target neurons, in the presence of voltage-gated Na^+^ channel blocker TTX (1 μM) and NBQX. Patch-clamp recordings were performed from PCs’ target neurons surrounded by many EGFP-positive PC axon terminals, which presumably correspond to neurons in DCN as suggested in a previous study (Kawaguchi and Sakaba, 2015). Consequently, a substantial population of observed mIPSCs was expected to be caused by spontaneous release from PC boutons compared to other inhibitory neurons (Figure 4A). Forskolin increased the frequency of mIPSCs significantly (13.0 ± 2.7 Hz to 19.0 ± 3.9 Hz, p = 0.013), without changes in the amplitude of mIPSCs (45.3 ± 12.6 pA to 39.9 ± 6.8 pA, p = 0.431; Figure 4B-D), in accord with previous studies on other neurons (Chen and Regehr, 1997; Kaneko and Takahashi, 2004). Thus, our results suggest that cAMP potentiates spontaneous release from PC axon terminals, as in other CNS neurons, although we cannot exclude the possibility that the forskolin-caused increase in mIPSC frequency was ascribed to augmented release from boutons of inhibitory neurons other than PCs. In addition, postsynaptic responsiveness is not affected by cAMP.

**Figure 4.**
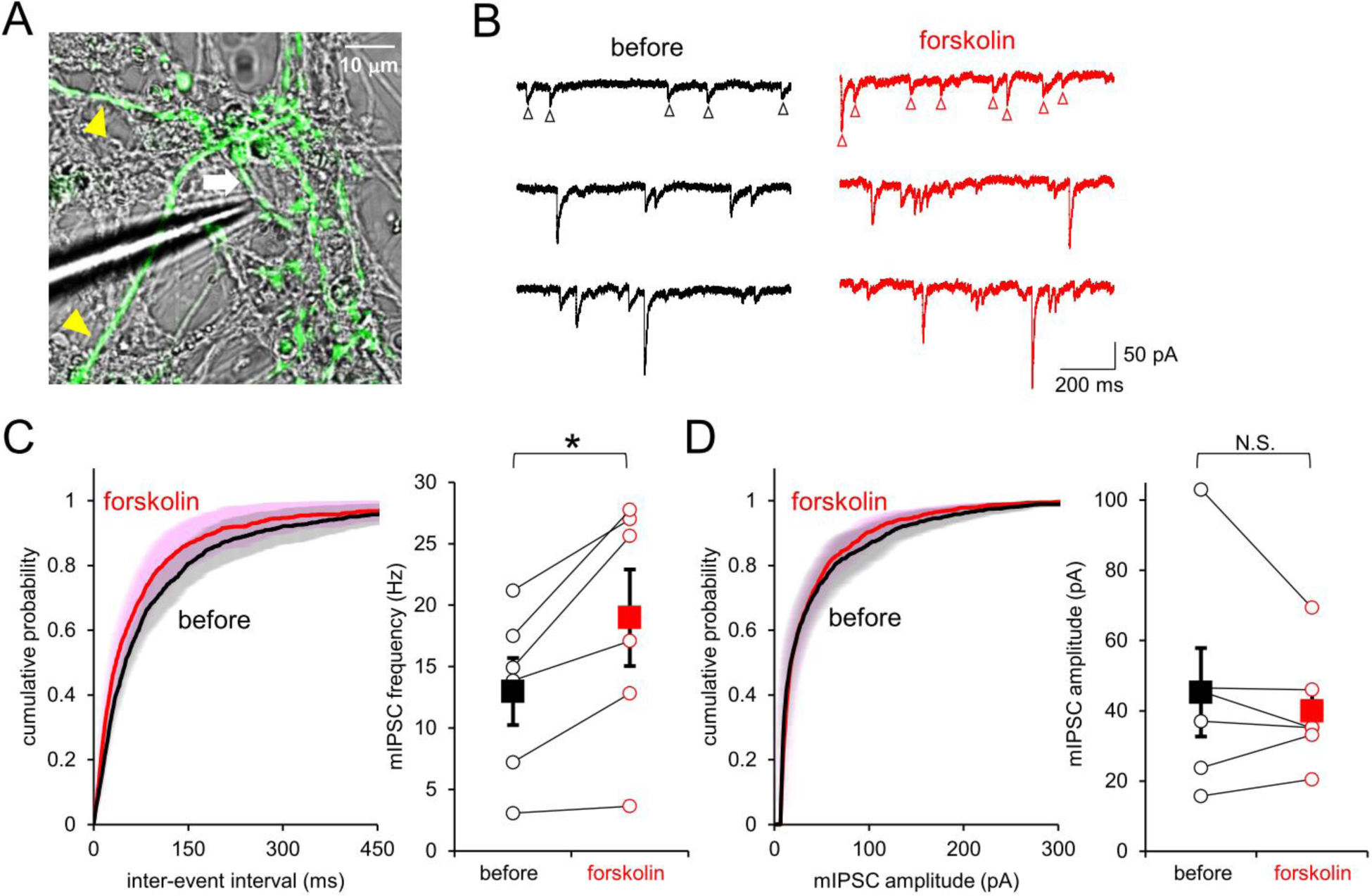
**A,** Image of a PCs’ target neuron (white arrow) under whole-cell recording surrounded by EGFP-positive PC boutons. **B**, Representative traces of mIPSCs (open arrowheads) before (left, black) and after (right, red) forskolin application. **C**, **D**, Left, Cumulative probability distribution of mIPSC inter-event intervals (C) and amplitudes (D) before (black) and after (red) forskolin application. SEM are also shown by thin colors. n = 6 cells. Right, Frequency (C) and amplitude (D) of mIPSC in individual cells (open circles), and mean ± SEM (closed squares) before and after application of forskolin. n = 6. Frequency, p = 0.013; Amplitude, p = 0.431

To directly obtain insights into the augmentation of the transmitter release by cAMP and how apparently counteracting actions of cAMP on presynaptic function, enhanced release probability and attenuated AP amplitude, totally control the release, we further performed paired patch-clamp recordings from a presynaptic PC terminal and the postsynaptic neuron (Figure 5A). We first examined whether cAMP modulates the AP-induced Ca^2+^ influx and/or the subsequent neurotransmitter release. For this end, two distinct AP waveforms, one in naïve condition and the other after forskolin application, were actually recorded in current clamp configuration from a PC terminal (Figure 5B), and those AP waveforms were applied to the voltage-clamped axon terminals to measure subsequent presynaptic Ca^2+^ currents. As shown in Figure 5C and 5E, presynaptic Ca^2+^ current was not affected by the application of cAMPS-Sp, suggesting that the Cav channels are insensitive to cAMP increases in a PC bouton, in accord with other neuronal presynaptic terminals (Kaneko and Takahashi, 2004; Midorikawa and Sakaba, 2017). In contrast to the presynaptic I_Ca_^2+^ insensitive to cAMP modulation, the postsynaptic target cell exhibited a tendency to increase in the postsynaptic conductance change by cAMP estimated from eIPSCs upon the identical presynaptic AP command, although not statistically significant (in average 218%; Figure 5D, F). The postsynaptic response relative to presynaptic I_Ca_^2+^ showed ∼ 2.1 fold higher efficiency in the presence of cAMPS-Sp (p = 0.028, Student’s t-test; Figure 5G).

**Figure 5.**
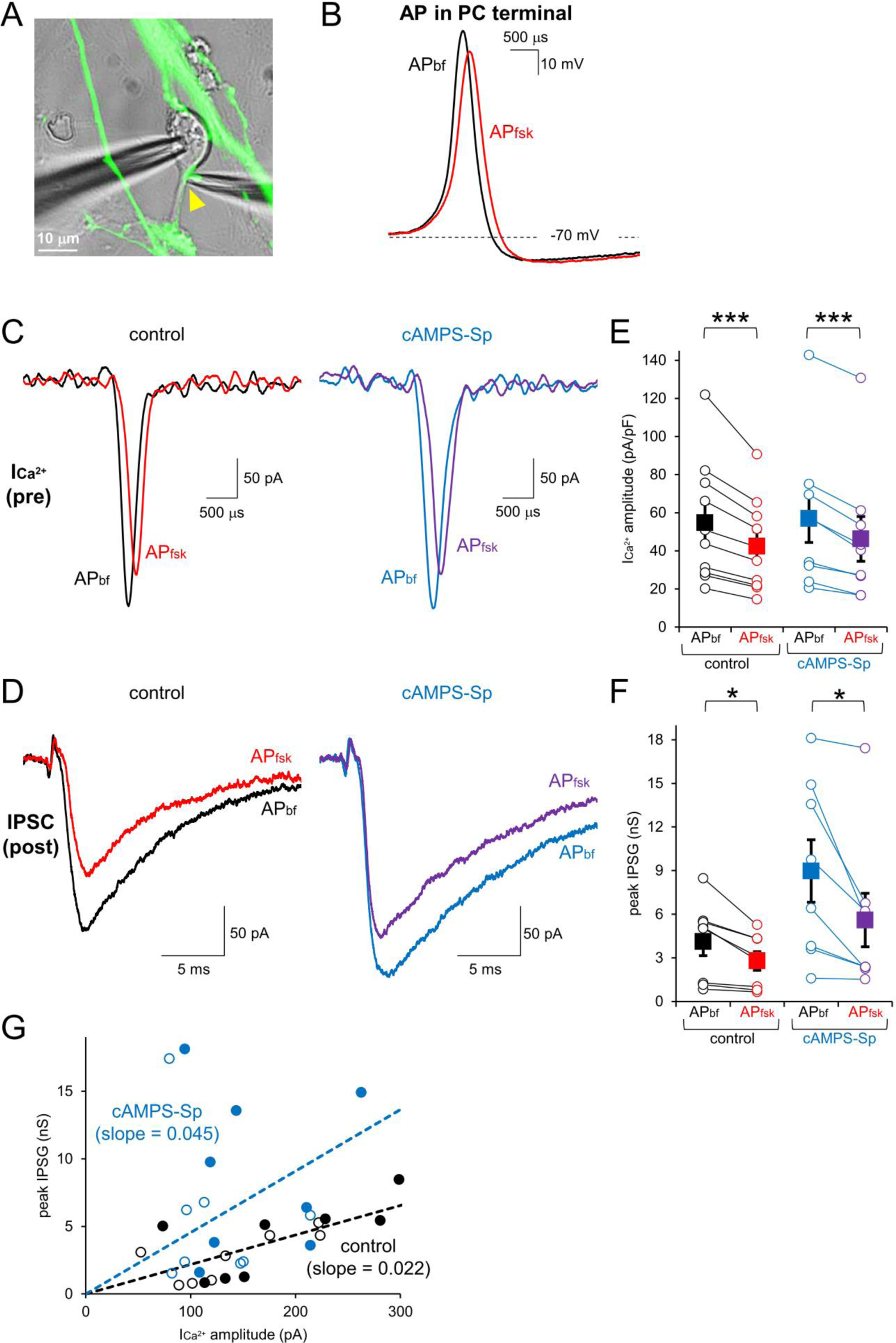
**A,** Image of paired recordings from a presynaptic PC bouton (yellow arrowhead) and its postsynaptic target neuron. **B**, AP waveforms recorded from a PC terminal before (black, AP_bf_) and after (red, AP_fsk_) bath forskolin application. **C**, **D**, Representative presynaptic Ca^2+^ currents (C) and IPSCs (D) in response to presynaptic voltage command of AP_bf_ or AP_fsk_ without (control, left) or with cAMPS-Sp in the internal solution (right). **E**, **F**, Average amplitudes of Ca^2+^ currents (E) and inhibitory postsynaptic conductances (IPSG, F) of individual cells (open circles), and mean ± SEM (closed squares) with or without cAMPS-Sp. For I_Ca_^2+^, data are normalized by the size of presynaptic membrane capacitance under the voltage-clamp. **G**, Relationship between IPSG and Ca^2+^ current upon AP_bf_ (filled circles) or AP_fsk_ (open circles) with or without cAMPS-Sp. Linear fits to the data for control and cAMPS-Sp, exhibiting higher slope for the latter.

On the other hand, the weakened AP waveform obtained after the forskolin application (as shown in Figure 5B) decreased the amplitude of presynaptic Ca^2+^ current to 77 ± 1% (control) and 79 ± 2% (cAMPS-Sp) compared with that triggered by the original AP waveform (Figure 5C, E). As a result of reduction of presynaptic I_Ca_^2+^, the postsynaptic conductance change estimated from eIPSCs also exhibited a reduction in amplitude to 71 ± 3% (control) and 64 ± 8% (cAMPS-Sp) (Figure 5D, F). Interestingly, the size of postsynaptic responses upon a smaller AP waveform in the presence of cytoplasmic cAMPS-Sp was similar to that upon the original AP waveform in the control condition (Figure 5D, F). All these results taken together suggest that the cytoplasmic cAMP increase might augment release of synaptic vesicles without affecting presynaptic Ca^2+^ channels, and that the cAMP-mediated AP attenuation accompanied with a decrease of presynaptic Ca^2+^ influx would counteract the cAMP-mediated augmentation of release, resulting in an apparent constancy of synaptic output size in PCs (as observed in Figure 1).

We next attempted to study in detail the cAMP-caused augmentation of transmitter release in a PC bouton, where transmitter release machinery is thought to be tightly coupled to Ca^2+^ channels (Diaz-Rojas et al., 2015), unlike glutamatergic synapses characterized by low release probability and cAMP-mediated facilitation via shortening of the Cav-release coupling distance, such as hippocampal mossy fibers boutons (Midorikawa and Sakaba, 2017). For that aim, we performed direct voltage-clamp recordings from a PC axon terminal and applied square pulse depolarization (to 0 mV for 1 – 50 ms duration), and recorded Ca^2+^ currents and the resultant changes of presynaptic membrane capacitance (Cm) which reflects total amount of exocytosed synaptic vesicles. In line with the lack of effect of cAMP on Ca^2+^ channels (as shown in Figure 5), the Ca^2+^ current in response to square depolarization was not significantly altered by cAMPS-Sp (Figure 6A, C). On the other hand, the Cm increased more efficiently even upon shorter depolarization pulses in the presence of cAMPS-Sp (11.7 ± 2.8 to 20.5 ± 2.4 fF/pF upon 5 ms pulse, p = 0.028), whereas upon longer pulses reaching the maximal value of Cm increase similar to the control condition (28.5 ± 8.2 to 29.0 ± 7.4 fF/pF upon 50 ms pulse, p = 0.97, see Figure 6B, D). Thus, the increase in the cytoplasmic cAMP seems not to change the total size of readily releasable pool of synaptic vesicles, but to augment the efficiency of Ca^2+^ to trigger the exocytosis of vesicles. These results demonstrated that even a ‘tight coupling’ presynaptic terminal of a PC exhibits cAMP-caused facilitation of transmitter release through increased release probability, in a similar manner to other synapses (Yao and Sakaba, 2010).

**Figure 6.**
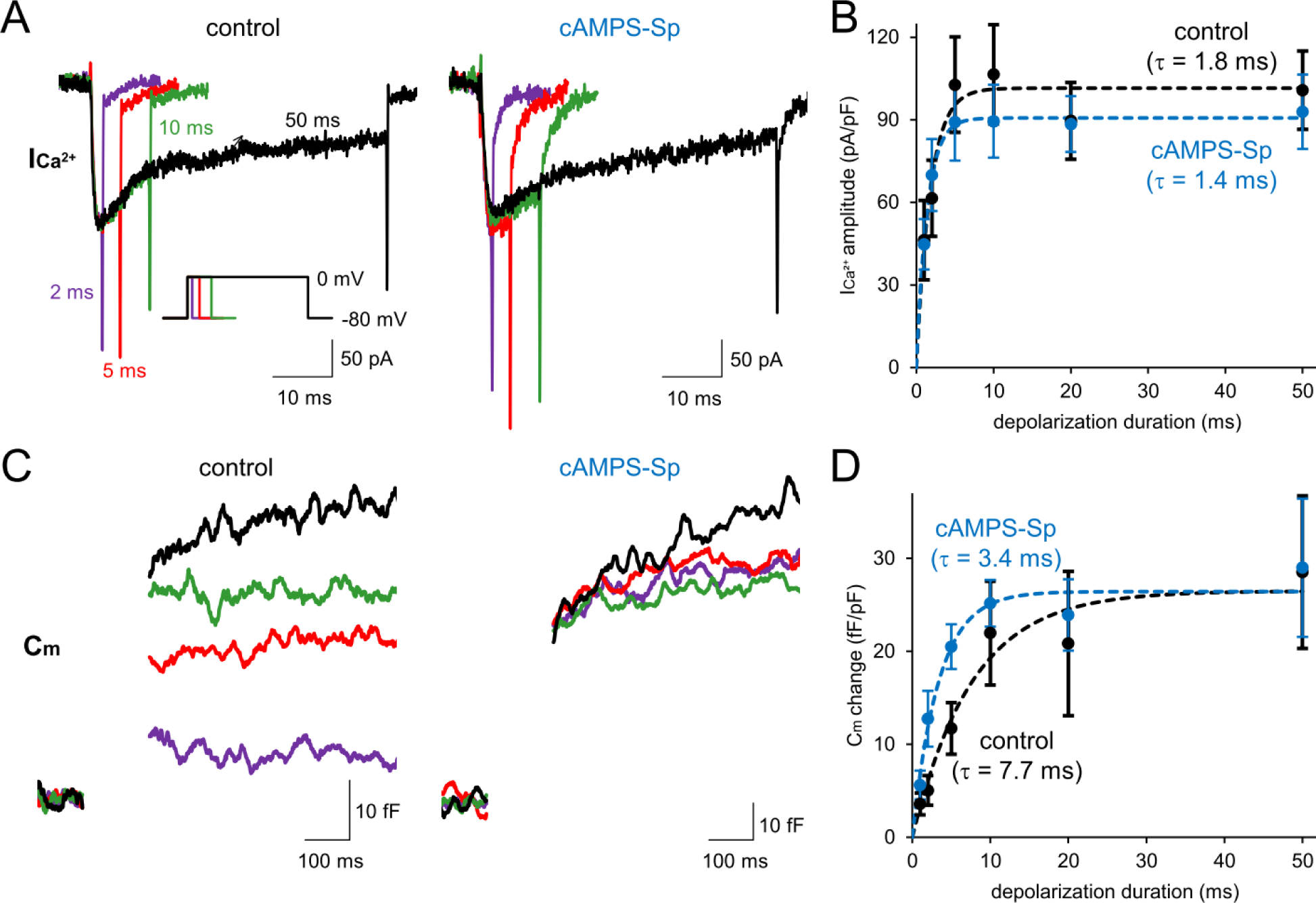
**A, B**, Representative traces of presynaptic Ca^2+^ currents (A) and the resultant Cm jump (B) upon 2, 5, 10 and 50 ms of depolarization (inset) without (left) or with cAMPS-Sp (right). **C**, **D**, Ca^2+^ current amplitudes (C) and Cm changes (D) upon depolarization pulses without or with cAMPS-Sp. Data are represented as mean ± SEM. Single exponential fits in each condition are shown by dotted lines. I_Ca_^2+^, n = 11 (control) and 12 (cAMPS-Sp) boutons, p = 0.154; Cm changes, n = 11 (control) and 12 (cAMPS-Sp) boutons, p = 0.102.

### Selective slowing of AP conduction by cAMP at a PC axon in acute slice

We finally attempted to confirm the cAMP’s action on PC output synapses also in DCN of acute slices prepared from cerebellar vermis at P12-20 rats. Injection of an AAV vector, AAV2-CA-EGFP, into the cerebellum of P2-5 rat pups fluorescently labeled PC axons a few weeks later (Figure 7A). To examine whether cAMP slows down the AP conduction in a PC axon in slice, we performed a direct patch-clamp recording from an axonal bleb, an enlarged structure formed at the cut end of the axon during slice preparation. APs were elicited by extracellular electrical stimulation with a glass pipette placed at > 200 μm away from the recording site (Figure 7A). We measured delays of the AP arrival from the stimulation site (Figure 7B). Application of forskolin increased the delay significantly (Figure 7B, C). Although we cannot completely exclude the possibility that forskolin slowed down the AP initiation upon the stimulation rather than the conduction, together with the paired soma-axon recordings in culture (as shown in Figure 2), our results indicate that cAMP decreases the AP conduction velocity in a PC.

**Figure 7.**
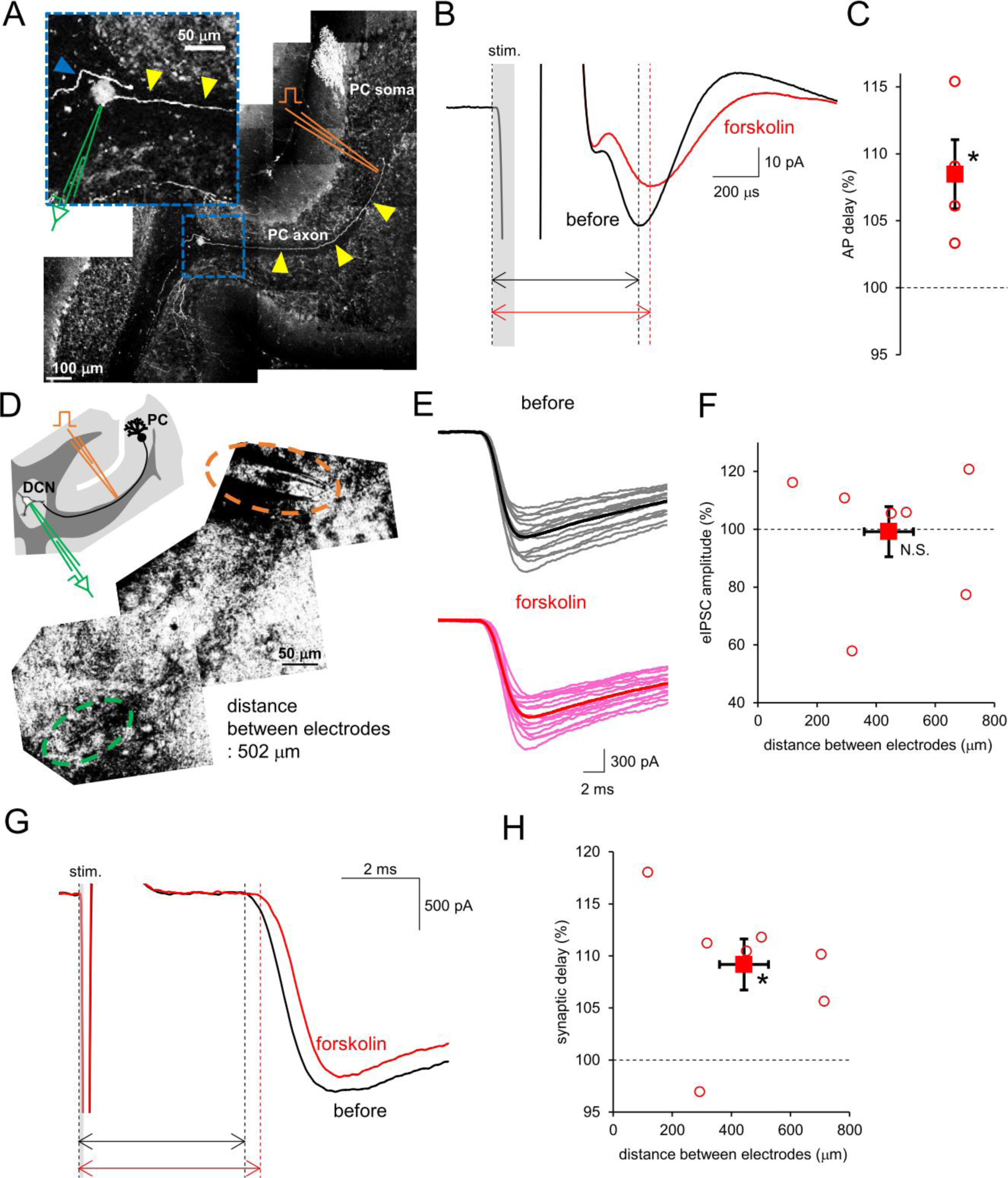
**A,** Images of EGFP-labeled PC axons (yellow arrowheads; blue indicates another one) in a cerebellar acute slice. Magnified image of the recorded axonal bleb is shown in inset (green, schematical electrode). A pipette for electrical stimulation (orange) was placed at ∼ 950 μm away from the recorded bleb. **B**, **C**, Representative averaged traces (B) and relative delay (C, data for individual cells and mean ± SEM) of cell-attached axonal APs before (black) and after (red) forskolin application. Delays of cell-attached AP peak from the stimulation onset were measured. n = 4, p = 0.043. **D**, Experimental scheme of eIPSC recordings from a DCN neuron. Electrical stimulations at white matter elicited APs in PC axons. **E**, **F**, Representative traces (E) and normalized average amplitudes (F) of eIPSCs after forskolin application. Individual (gray, pink) and average (black, red) traces are shown. n = 7, p = 0.655. **G**, **H**, Representative traces (G) and normalized synaptic delays (H) of eIPSCs before (black) and after (red) forskolin application. n = 7, p = 0.021.

We then examined whether the evoked transmitter release is altered by cAMP at PC-DCN synapses in slice. Whole-cell patch-clamp recording was performed from a neuron with a large cell body (> 15 μm diameter) in the DCN region. eIPSCs were triggered by electrical stimulation of the white matter in the presence of NBQX (Figure 7D, E). In accord with previous studies, eIPSCs from PC axon terminals were identified by almost all-or-none responses with large amplitude upon elevating stimulation intensity (Figure 7E, Telgkamp and Raman, 2002). Forskolin did not significantly change the amplitude of eIPSCs, although variant (Figure 7E, F). Neither eIPSC time courses nor CV of amplitude was affected by forskolin (Supplementary Figure 6). On the other hand, as in primary culture, forskolin significantly increased the synaptic delay (Figure 7G, H). Taken these results in slices together, AP is specifically slowed down in conduction by cAMP during axonal propagation without clear change of output sizes, in consistence with results obtained in a primary culture.

## Discussion

In this study, taking advantages of direct patch-clamp recordings from an intact long axon of PC in culture and from an axonal cut end in acute slice, we found that AP is reduced in size by increase of cytoplasmic cAMP specifically in an axon through reduction of axonal Na^+^ currents, which results in slower conduction of APs along an axon and hence longer delay in synaptic outputs. In parallel, Ca^2+^ influx at a presynaptic terminal is also negatively regulated indirectly by cAMP via weakened size of AP that counteracts the facilitatory effect of cAMP on transmitter release machinery. As a result of interplay of intricate mechanisms, cAMP elevation in a PC axon specifically delays the speed of axonal signaling and hence output to the target neurons, keeping almost constant synaptic output strength. Thus, our present study demonstrated a mechanism to separately modulate the temporal precision of axonal signal processing independent of the signal intensity.

### Analogue modulation of axonal signaling

A large body of previous works revealed that the presynaptic AP waveform is a critical determinant of synaptic strength because it sets the maximal probability and time course of Ca_v_ opening (Borst and Sakmann, 1999; Geiger and Jonas, 2000; Kawaguchi and Sakaba, 2015; Sabatini and Regehr, 1997; Taschenberger and von Gersdorff, 2000; Zbili et al., 2020). Direct patch-clamp recordings from PC axonal trunk/boutons here demonstrated that cAMP elevation significantly reduces amplitudes and slows down time courses of APs specifically at axon, lowering the AP conduction velocity. This contrasts to the classical view of axonal AP as an all-or-none high fidelity signal, and suggests that any changes in waveforms might also change the speed of axonal signaling. Previous studies showed that axonal AP waveform changes upon high frequency firing in axons of a variety of neurons, such as cerebellar PCs (Kawaguchi and Sakaba, 2015), hippocampal mossy fibers (Alle and Geiger, 2006; Engel and Jonas, 2005), fast spiking interneurons in hippocampus (Hu and Jonas, 2014), and Calyx of Held in the auditory pathway of brain stem (Taschenberger and von Gersdorff, 2000). It would be interesting to look at the constancy of output timing during repetitive firing in those neurons. In addition, as a result of altered AP waveform, Ca_v_ is differentially activated and transmission altered (Borst and Sakmann, 1999; Carta et al., 2014; Geiger and Jonas, 2000; Kawaguchi and Sakaba, 2015; Kawaguchi and Sakaba, 2017; Sabatini and Regehr, 1997). On the other hand, recent direct axonal patch-clamp recordings suggest that other neuronal axons, such as fast spiking GABAergic interneurons in the cerebellar molecular layer or in the layer 5 of cerebral cortex, exhibit very stable AP waveforms even at very high frequency firing (Ritzau-Jost et al. 2021; Trigo and Kawaguchi, 2023), although modulation of AP width by inactivation of Kv channels was also reported (Rowan and Christie, 2017). Furthermore, axonal AP waveforms have been shown to be affected by local receptors for various transmitters (Trigo et al., 2010; Zorrilla de San Martin et al., 2017), such as GABA.

Our paired recordings from the soma and axonal trunk yield the AP conduction velocity in a PC axon as ∼ 540 μm/ms at room temperature (see Figure 1), which compromises the velocity around 770 μm/ms studied in slice at physiological temperature (Clark et al., 2005), when considering the temperature sensitivity of AP conduction as 1.5 ∼ 2 fold per 10 °C changes (Q_10_) (Chapman, 1967). The AP conduction velocity was here slowed down as a result of altered AP waveforms by cAMP through lowered membrane excitability, leading to delayed synaptic outputs from terminals. On the other hand, increased membrane excitability by cAMP through activating HCN channels was also shown in previous studies to change the axonal AP conduction velocity in cerebellar and cortical neurons, although the direction of speed change was diverse: speeding in cerebellar parallel fibers and mossy fibers (Byczkowicz et al., 2019), whereas slowing in cortical pyramidal cells (Lezmy et al., 2021). In addition, the decay phase of presynaptic AP by itself also affects the synaptic latency by changing the timing of Ca^2+^ influx, as shown in cortical synapses between layer 5 neurons (Boudkkazi et al., 2011). Thus, the timing of neuronal outputs are fine-tuned by dynamic interplay of multiple mechanisms to determine the AP waveforms, which would be central for neuronal signal integration.

### Mechanism of AP modulation and its impact on release

Our voltage-clamp recordings from axonal trunk and boutons of PCs clarified negative regulation of Na^+^ channels by cAMP specifically at axon (see Figure 3), which gives rise to the attenuated AP. As far as we know, this is the first demonstration of clear decrease of AP sizes by quick modulation of Na^+^ channels through intracellular signaling pathway. In avian auditory system, Kuba and colleagues have shown that Nav channels are regulated in density and position around the axonal initial segment in a manner dependent on neuronal activity, resulting in altered threshold of axonal firing (Kuba et al., 2010). Taking into consideration that Na^+^ channels are the primary factor producing excitatory drive for membrane, control of Na^+^ channels at axon would have a powerful effect on axonal signaling in general, as shown here. PCs are reported to express various types of Na^+^ channels, such as Nav1.1, 1.4, 1.5, 1.6, 1.8 and 1.9 (Schaller and Caldwell, 2003; Allen Brain Atlas), and it remains unclear what type of Nav is responsible for axonal component susceptible for the down-regulation by cAMP and hence attenuated APs, which should be clarified in future studies. The fact that the cAMP-caused reduction of Na^+^ current was specifically evident in axon, but not at the soma, implies different subtypes of Nav located at the soma and axon, which makes it feasible to specifically control the axonal AP waveform. Our data suggest that downstream molecule of cAMP other than PKA or EPAC underlies the negative regulation of Nav, which remains to be clarified. One of such cAMP effectors, named POPCDs, might play a role in the Nav modulation. On the other hand, in contrast to previous studies showing cAMP-caused positive modulation of hyperpolarization-activated HCN currents in other neurons (Byczkowicz et al., 2019; Lezmy et al., 2021), PC axons showed no clear change of HCN currents upon cAMP elevation (see Figure 3C, D), in line with previous reports of selective expression of cAMP-insensitive HCN1 in PCs (Notomi and Shigemoto, 2004; Wang et al., 2001). Thus, different neuronal axons might have distinct types of channels, such as Nav and HCN, and signaling molecules, giving rise to diverse mechanisms to change the AP waveform and conduction velocity. Other than regulation of Nav, numbers of previous studies have provided evidences for changes of AP waveforms through modulation of K^+^ channels in various types of neuronal axon, such as those in hippocampal pyramidal cells (Cho et al., 2020; Hoppa et al., 2014), and molecular layer interneuron (Begum et al., 2016; Rowan and Christie, 2017).

AP waveforms powerfully control the presynaptic Ca^2+^ influx, thereby impacting on the transmission, although the extent is variable depending on the neuronal type (Boudkkazi et al., 2011). When the opening kinetics of presynaptic Ca^2+^ channels is fast enough relative to the AP waveforms, a majority of Ca^2+^ channels has a chance to get activated upon a single AP even if the peak amplitude is somehow altered (Bischofberger et al., 2002; Borst and Sakmann, 1998). In such a case, Ca^2+^ influx is predominantly determined by how the depolarized potential decays, during which Ca^2+^ is allowed to gradually enter as the driving force gets larger. Indeed, presynaptic boutons of Calyx of Held synapses or hippocampal mossy fibers exhibit more powerful effects of the time course rather than amplitude of APs on the presynaptic Ca^2+^ influx and the resultant transmitter release (Bischofberger et al., 2002; Geiger and Jonas, 2000). On the other hand, only a limited population of Ca^2+^ channels open in the cases when AP at a bouton is rapid compared to the kinetics of Ca^2+^ channel activation, giving rise to larger sensitivity of Ca^2+^ influx to the AP amplitude. Indeed, changes of AP amplitude have been shown to control the Ca^2+^ influx in boutons of cerebellar neurons like PCs, granule cells, and GABAergic interneurons (Kawaguchi and Sakaba, 2015; Kawaguchi and Sakaba, 2017; Trigo and Kawaguchi, 2023). Particularly at synapses which are functionally designed to undergo transmitter release based on tight coupling between Ca^2+^ channels and release machinery, the number of Ca^2+^ channels activated, rather than the total amount of Ca^2+^ entering into the cytoplasm, mainly determines the amount of vesicles undergoing exocytosis. Thus, AP waveform and the resultant Ca^2+^ influx dynamically interact to differently control synaptic outputs depending on elaborate presynaptic design, which would decide whether and to what extent the AP modulation, as shown here by cAMP, impacts the strength of synaptic outputs.

### Release machinery modulated by cAMP

One of the major second messenger cAMP, has been shown to modulate synaptic transmission working not only on postsynaptic structure (Blitzer et al., 1995), but also on presynaptic boutons at wide variety of neurons (Byrne and Kandel, 1996; Capogna et al., 1995; Chavez-Noriega and Stevens, 1994; Chen and Regehr, 1997; Cheung et al., 2006; Huang et al., 1994; Kaneko and Takahashi, 2004; Meadows et al., 2021; Saitow et al., 2000; Salin et al., 1996; Trudeau et al., 1996; Weisskopf et al., 1994). In addition, cAMP is also known to be involved in long-term potentiation via augmented transmitter release at low release probability synapses, such as hippocampal mossy fibers (Kaeser-Woo et al., 2013; Midorikawa and Sakaba, 2017; Nicoll and Malenka, 1995) and cerebellar parallel fibers (Salin et al., 1996). Thus, the mechanism how cAMP augments the transmitter release has attracted attention. Recent elegant studies applying direct patch-clamp recordings combined with fluorescent imaging of vesicle fusion to the plasma membrane and super-resolution fluorescent microscopy on hippocampal mossy fibers demonstrated that cAMP shortens the coupling distance between Ca^2+^ channels and release machinery (Fukaya et al., 2021, 2023). This shift of Ca^2+^ channel coupling to the release machinery from loose to tight one is thought to be reflected as the increased release probability, as experimentally characterized by the higher resistance of transmission to slow Ca^2+^ chelator EGTA in the cytoplasm. Although the shortening of the distance of Ca^2+^ channels to release machinery could be an efficient way to positively modulate the transmission, it is unlikely that such a mechanism strongly operates at tight-coupling synapses because of the limited margin for further shortening of the coupling distance. As shown in this study, PC boutons which are EGTA-resistant tight Cav-release coupling (Diaz-Rojas et al., 2015), undergo cAMP-mediated increase in release probability, as characterized by more efficient vesicular fusion upon smaller Ca^2+^ influx with changing neither Ca^2+^ influx (see Figure 5 and 6) nor total amount of release-competent vesicles (see Figure 6). Relatively weak facilitation of release by cAMP in PC boutons (∼ 2 fold) compared to strong mossy fiber boutons (∼ 4 fold) (Weisskopf et al., 1994) might be ascribed to the distinct synaptic design in terms of Cav-release coupling. Previous studies using paired pre- and postsynaptic recordings together with Ca^2+^ uncaging methods at Calyx of Held synapses clarified that cAMP potentiates transmitter release by increasing Ca^2+^ sensitivity of release machinery rather than by changing either the coupling distance or readily releasable pool size (Yao and Sakaba, 2010), in a similar manner to PCs as shown here. Recent studies are shedding light on critical roles of PKA and/or EPAC-mediated modulation of RIM1 and Munc-13 at active zones in the cAMP-mediated enhanced release (Fukaya et al., 2023; Wang et al., 2023). Similar molecular mechanisms might underlie also at the tight-coupling PC boutons, which is an important issue to be addressed in future studies.

## Materials and Methods

All experimental procedures were performed in accordance with regulations on animal experimentation in Kyoto University and approved by the local committee in Graduate School of Science, Kyoto University.

### Cerebellar primary cultures

The method for preparing primary dissociated cultures of cerebellar neurons was similar to that in a previous study (Kawaguchi and Sakaba, 2015). PCs were transfected with EGFP at 4-5 days after culture with an AAV vector serotype 2 under the control of CA promoter (AAV2-CA-EGFP). PCs could be visually identified by large cell bodies and thick dendrites. An axon was clearly distinct from dendrites which have high density of spines in a PC. Experiments were performed >25 days after preparation of the culture.

### Cerebellar slices

Acute sagittal slices (200 μm thickness) of cerebellar vermis were prepared from Wistar rats of either sex aged 12 to 16 days old for experiments on synaptic transmission, and 17 to 20 days old for those on AP conduction. After decapitation, the cerebellum was quickly removed and cut with a Leica vibroslicer (VT1200S) in an ice-cold sucrose solution containing the following (in mM): 60 NaCl, 120 Sucrose, 25 NaHCO_3_, 1.25 NaH_2_PO_4_, 2.5 KCl, 25 D-glucose, 0.4 ascorbic acid, 3 myo-inositol, 2 Na-pyruvate, 0.1 CaCl_2_, 3 MgCl_2_, pH 7.3-7.4, osmolarity 300-350 mOsm/KgH_2_O with continuous bubbling with mixed gas (95% O_2_ and 5% CO_2_). Slices were then incubated at 37 °C for 30 min to 1 h in an extracellular solution containing the following (in mM): 125 NaCl, 25 NaHCO_3_, 1.25 NaH_2_PO_4_, 2.5 KCl, 25 D-glucose, 0.4 ascorbic acid, 3 myo-inositol, 2 Na-pyruvate, 2 CaCl_2_, 1 MgCl_2_, pH 7.3-7.4, 300-320 mOsm/KgH_2_O. For experiments on AP conduction, PCs were EGFP-labeled by injection of AAV vector (AAV2-CA-EGFP) into the cerebellum of rat at postnatal days 2 to 5.

### Electrophysiology

Patch-clamp recording was performed with an amplifier (EPC10, HEKA, Germany) at room temperature (20-24 °C), in an extracellular solution mentioned above for slices, or that containing the following for culture (in mM): 145 NaCl, 10 HEPES, 10 Glucose, 2 CaCl_2_, 1 MgCl_2_, pH 7.3-7.4 adjusted by KOH and osmolarity 300-310 mOsm/KgH_2_O. In some experiments, 2,3-Dioxo-6-nitro-1,2,3,4-tetrahydrobenzo[f]quinoxaline-7-sulfonamide (NBQX, 10 μM), picrotoxin (50 μM), tetrodotoxin (TTX, 1 μM) were applied to the bath to inhibit glutamatergic EPSCs, GABAergic IPSCs, and APs, respectively. For cell-attached recordings in both culture and slices, patch pipettes were filled with extracellular solutions. For current-clamp recordings, K^+^-based internal solution with the following composition (in mM) was used: 155 K-gluconate, 7 KCl, 10 HEPES, 0.5 ethylene glycol bis (β-aminoethylether) N,N,N’,N’-tetraacetic acid (EGTA), 2 Mg-ATP, 0.2 Na-GTP, pH 7.3-7.4 adjusted by KOH and osmolarity 320-330 mOsm/KgH_2_O. For voltage-clamp recordings from the axon terminal of a PC, a patch pipette was filled with a CsCl-based internal solution (pH 7.3-7.4 adjusted by CsOH and osmolarity 320-330 mOsm/KgH_2_O) containing (in mM): 170 CsCl, 10 HEPES, 0.5 EGTA, 2 Mg-ATP, 0.2 Na-GTP, and recordings were performed in the presence of external TTX and tetraethylammonium (TEA, 2 mM). For postsynaptic recordings from PC-target neurons, the internal solution was mixture of CsCl and Cs-gluconate-based solutions (pH 7.3-7.4 adjusted by CsOH and osmolarity 320-330 mOsm/KgH_2_O) containing (in mM): 137 or 23 CsCl, 33 or 147 Cs-gluconate, 10 HEPES, 5 EGTA, 2 Mg-ATP, 0.2 Na-GTP. In measurements of voltage-gated currents for Na^+^, K^+^ and HCN channels, K^+^-based internal solution was used. When gluconate-based internal solution was used, the liquid junction potential (∼ 15 mV) was corrected. H89 (1 μM) or ESI-09 (100 μM) was applied to the bath or the internal solution filling the patch pipette to inhibit PKA or Epac, respectively. To increase internal cAMP, forskolin (50 μM) or cAMPS-Sp (1 mM) was added extracellularly to the bath or intracellularly through the patch pipette, respectively. In some cases, experiments with dimethyl sulfoxide (DMSO, 0.05%) applied to the bath were used as control. The membrane potential of a PC was held at -70 mV unless otherwise stated. PCs’ target postsynaptic neurons in either culture or slice were voltage-clamped at -70 to -100 mV for IPSC recordings to avoid unclamped Na^+^ currents. Series resistance of the PC soma, axon and terminal was, 13 ± 5 MΩ, 65 ± 18 MΩ and 124 ± 39 MΩ (mean ± SD; n = 28, 44, 38 cells), corrected by 40-60%, respectively. On-line series resistance compensation (20-60%) was applied for IPSC recordings, and the remaining resistance was corrected off-line after the recording. eIPSCs in culture were accepted for analysis unless aberrant change in failure rate during recordings was observed.

APs were elicited by current injection of 500-900 pA for 10 ms into PC soma in culture or by extracellular electrical stimulations of ∼ 50 V or ∼ 5 mA for 10 or 100 μs with a glass pipette on the white matter in slices. In the recordings of AP conduction in slices, a glass pipette was placed at > 200 μm away from the recorded axonal bleb. To exclude the possible AP modulation at high frequency firing, axonal AP with intervals > 50 ms were used for analysis.

The onset of eIPSC was defined as the first time point at which the recorded current value showed change larger than 2SD of that at the basal condition. For mIPSCs analysis, individual cumulative probability of inter-event intervals and amplitudes, average amplitude, and frequency were calculated from *≥* 200 events which were *≥*7 pA with appropriate time courses which was visually confirmed.

The current (I_Na_^+^)-voltage (V_m_) relation for axonal voltage-gated Na^+^ currents (see Supplementary Figure 5) was fitted by the follow equation based on the Boltzmann function:

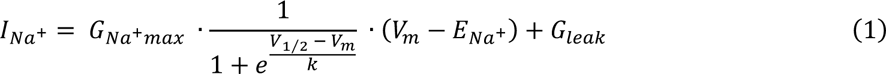

where E_Na_^+^ is the equilibrium potential of Na^+^. Fitting of I-V relationship for each axon yielded three key parameters for Nav activation, relative peak conductance of Na^+^ (G_Na_^+^_max_), voltage for half-maximal activation (V_1/2_), and the slope factor (k). In some experiments, postsynaptic responses were evaluated by calculating inhibitory postsynaptic conductance (IPSG) by dividing the IPSC amplitude by the driving force for Cl^-^.

Measuring of membrane capacitance (C_m_) was done using +DC technique (Neher and Marty, 1982) implemented on the Patchmaster software (HEKA, Germany). Presynaptic terminals were held at -80 mV and the sine wave (1 kHz and the peak amplitude of 30 mV) were applied on the holding potential. Because membrane conductance fluctuates during the depolarizing pulse, capacitance was usually measured between 200-300 ms after the depolarization.

To estimate the clamped area in a direct bouton recording, capacitive transients in response to a hyperpolarizing pulse (10 or 20 mV) were used. Capacitive transients at a terminal followed a single exponential with time constants of 0.39 ± 0.11 ms (mean ± SD, n = 37 cells), so that the clamped terminal size was estimated to be 3.6 ± 1.7 pF. Correlation coefficients between the clamping size and the amplitude of Ca^2+^ currents or Cm increase upon depolarization pulses for 50 ms (Figure 6) were 0.99, 0.59, respectively. To normalize the variation in Ca^2+^ currents and neurotransmitter release depending on the size of bouton, presynaptic Ca^2+^ current was normalized to the clamped membrane capacitance in each bouton (in Figure 5 and 6). In Figure 5, calculated IPSGs from postsynaptic neuron were also divided by presynaptic bouton size to normalize the amount of the release between terminals, although we cannot exclude the possibility that the postsynaptic conductance in response to the same presynaptic release of GABA is diverse between types of postsynaptic neurons.

### Statistics

Data are presented as mean ± SEM unless otherwise indicated. Statistical significance was evaluated by paired t-test, unpaired Students’ t-test, or two-way ANOVA.

## Acknowledgments

We thank Drs. Federico F. Trigo and Mitsuharu Midorikawa for the critical reading of the manuscript and helpful comments.

## Competing interests

The authors declare no competing interests.

## Funding

Japan Society for Promotion of Science, KAKENHI grants 22H02721 (SK), 22K19360 (SK), and 21K15189(TI).

Takeda Science Foundation (SK).

Naito Foundation (SK).

JST SPRING, Grant Number JPMJSP2110 (KF).

---

**Supplementary Figure 1.**
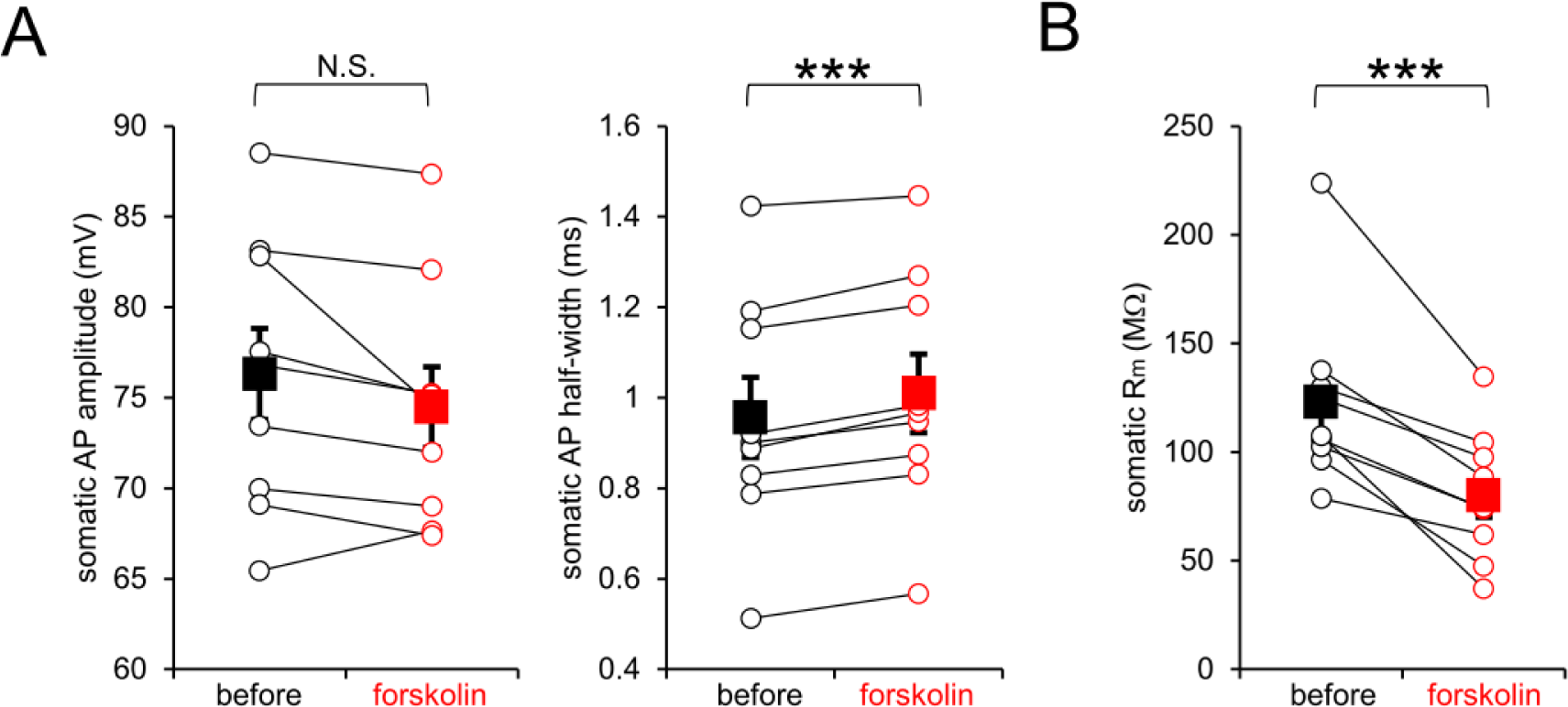
**A**, Average amplitudes (left) and halfwidths (right) of APs at PC soma of individual cells (open circles) and mean ± SEM (closed squares) before and after application of forskolin. n = 9; amplitude, p = 0.08; halfwidth, p = 0.000018 **B**, Apparent membrane resistances (Rm) at the soma of individual PCs (open circles) and mean ± SEM (closed squares) before and after (connected by lines) application of forskolin. n = 9, p = 0.000064

**Supplementary Figure 2.**
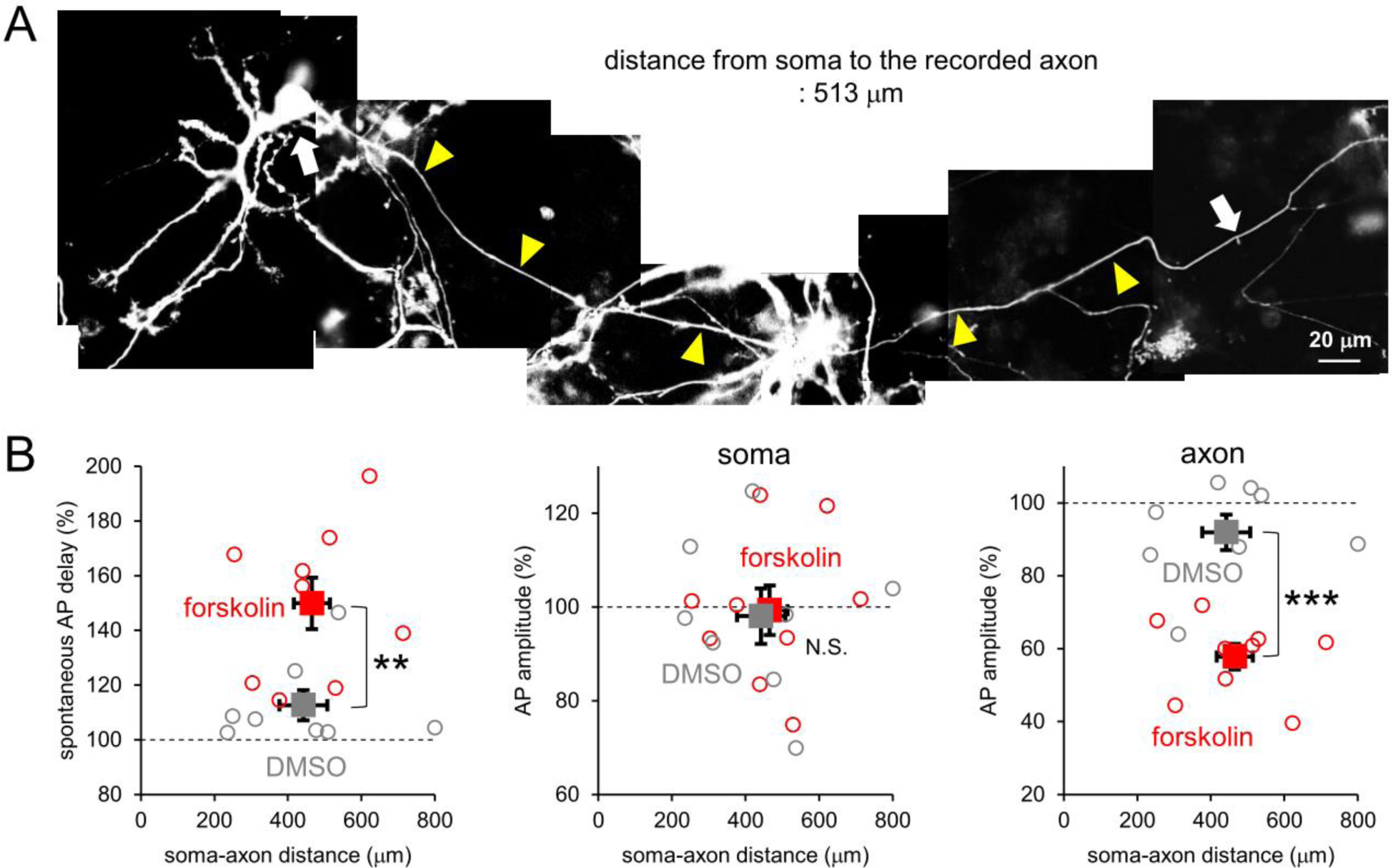
**A**, An Image of an EGFP-labeled cultured PC paired-recorded at the soma and the axonal site (white arrows). Magnified images of recording sites are shown in Figure 2A. Yellow arrowheads show the axon. The distance between two recorded sites along the axonal trajectory was 513 μm. **B**, Normalized averaged AP delays (left), and amplitudes at soma (middle) or axon (right) of individual cells (open circles) and mean ± SEM (closed squares) after application of DMSO (gray) or forskolin (red) plotted against the soma to axonal distances. Plots of delays for forskolin are the same as data in Figure 2C. n = 8 (DMSO) and 9 (forskolin). delay: p = 0.0049; amplitude: soma, p = 0.875; axon, p = 0.000036

**Supplementary Figure 3.**
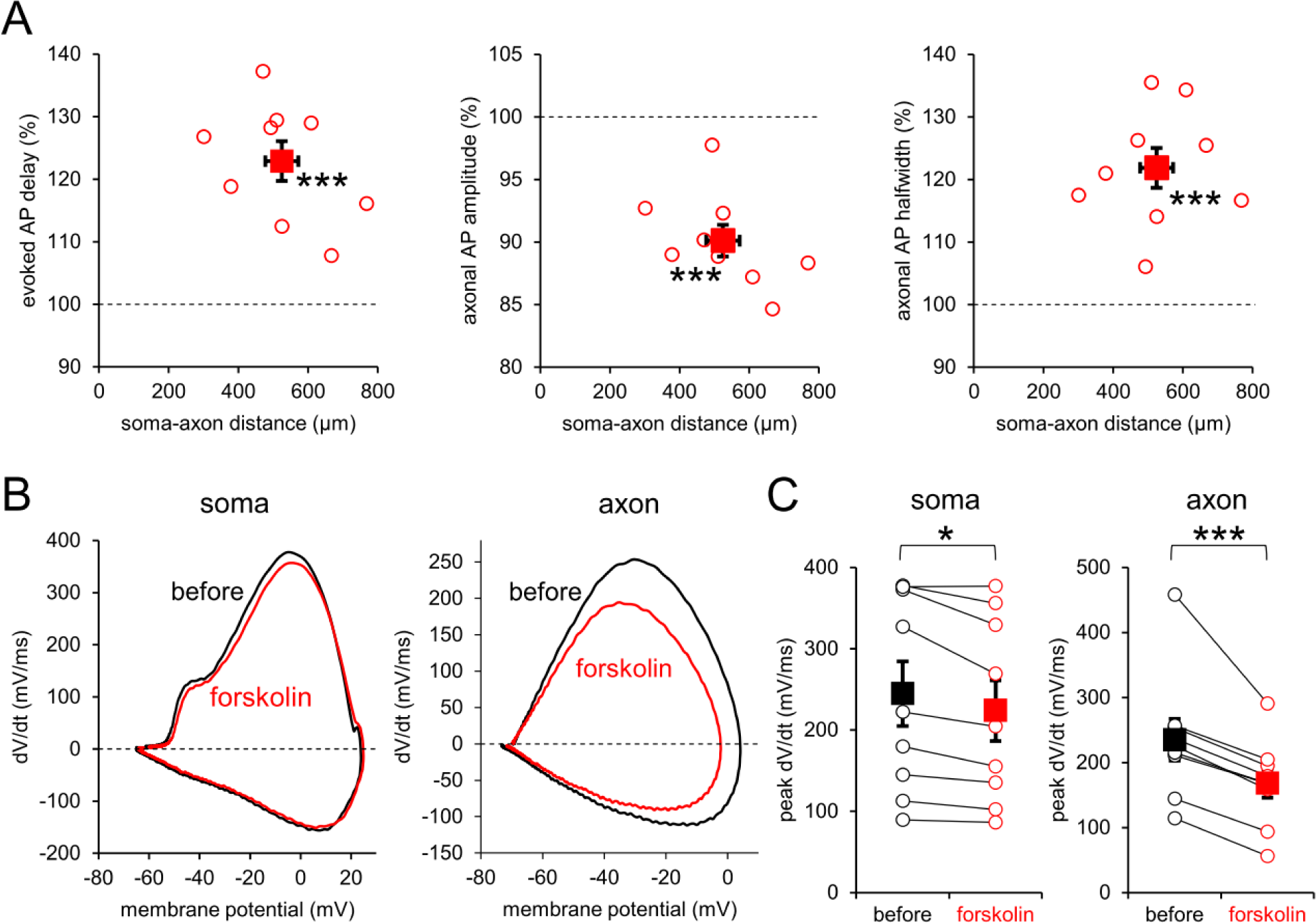
**A**, Normalized averaged AP delays (left), amplitudes (middle) and half-widths (right) at axon of individual cells (open circles) and mean ± SEM (closed squares) after forskolin application of plotted against the soma to axonal distances. n = 9; delay, p = 0.000033; amplitude, p = 0.000062; halfwidth, p = 0.00016 **B**, Phase plots of the APs (dV/dt vs V) at soma (left) and axon (right) (depicted in Figure 2D, E, respectively) before (black) and after (red) application of forskolin. **C**, The peak dV/dt of the APs at soma (left) and axon (right) of individual cells (open circles) and mean ± SEM (closed squares) before and after (connected by lines) forskolin application. n = 9; soma, p = 0.011; axon, p = 0.00079

**Supplementary Figure 4.**
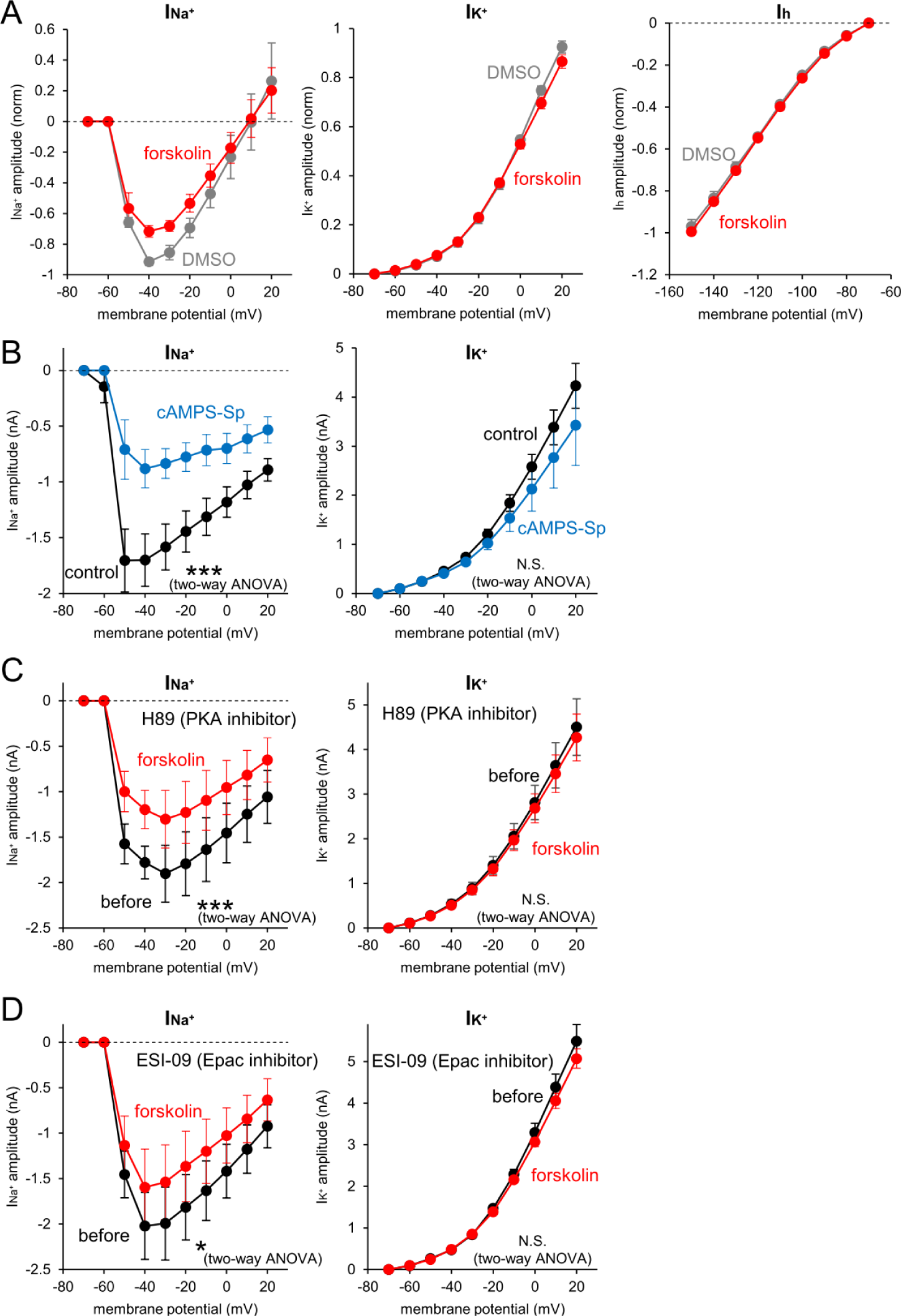
**A**, I-V relations of axonal voltage-dependent currents before (without) and after (with) forskolin (cAMP-Sp) voltage-gated I_Na_^+^ (left), I_K_^+^ (middle) and I_h_ (right) currents after application of DMSO (gray) or forskolin (red). Currents were normalized by the maximum I_Na_^+^ in each recorded axon before application. n = 6 (DMSO) and 7 (forskolin) axons. I_Na_^+^, p = 0.084; I_K_^+^, p = 0.130; I_h_, p =0.088 **B**, I-V curves of axonal voltage-gated I_Na_^+^ (left) and I_K_^+^ (right) without (control, black) or with cAMPS-Sp in the internal solution (blue). n = 6 (control) and 6 (cAMPS-Sp) axons. I_Na_^+^, p = 0.0000083; I_K_^+^, p = 0.506 **C**, **D**, I_Na_^+^ (left) and I_K_^+^ (right) before (black) and after (red) forskolin application in the presence of H89 (C) or ESI-09 (D) in the bath or internal solution, respectively. n = 6 (H89) and 6 (ESI-09) axons. I_Na_^+^: H89, p = 0.00048; ESI-09, p = 0.023; I_K_^+^: H89, p = 0.536; ESI-09, p = 0.078. Data are presented as mean ± SEM.

**Supplementary Figure 5.**
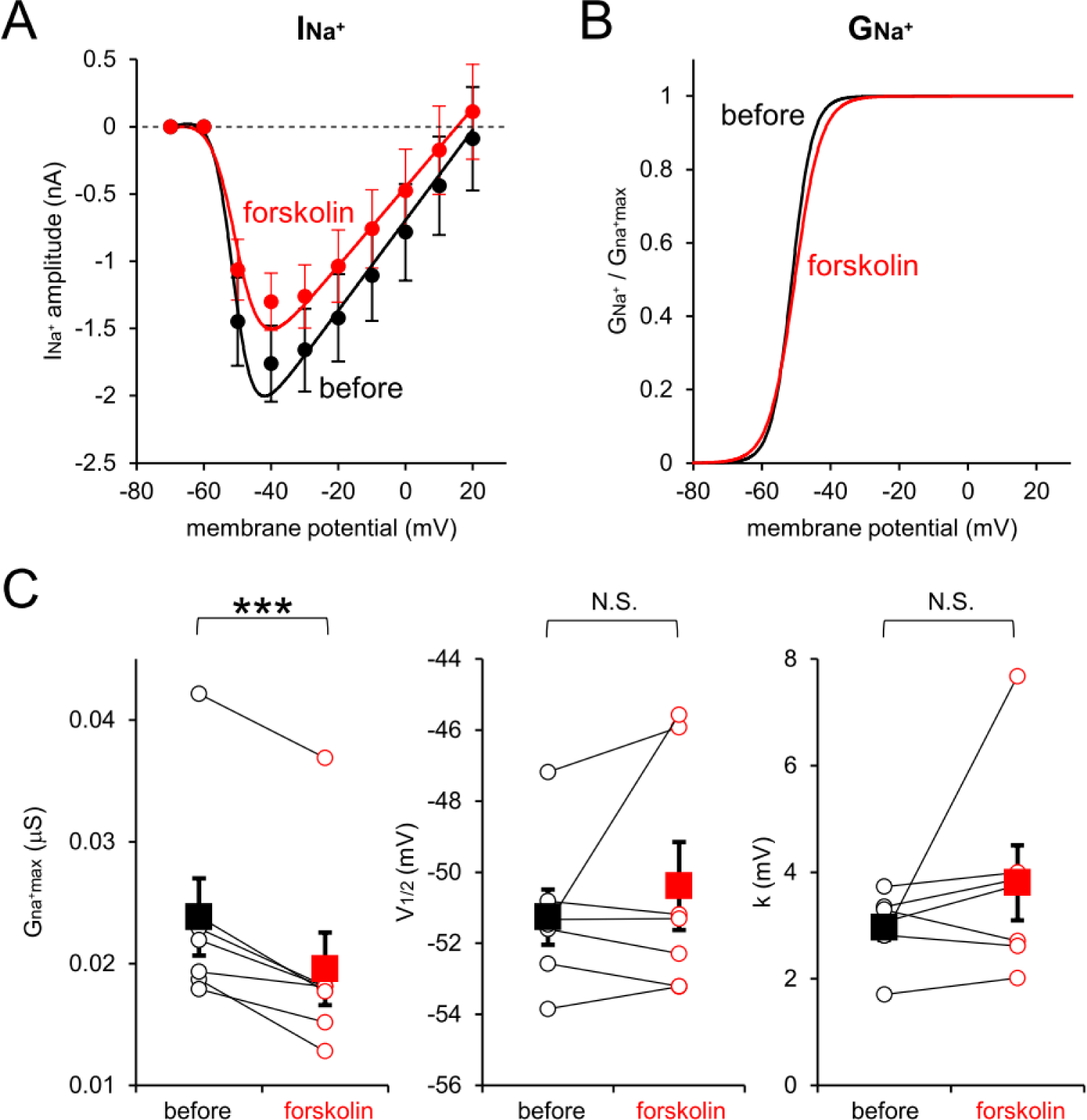
**A**, Fittings of the I-V curves for axonal voltage-gated I_Na_^+^ before (black) and after (red) application of forskolin (solid lines, see also **Materials and methods**). Plots are the same as data in Figure 3B. **B**, Activation curves of axonal voltage-gated Na^+^ conductance before (black) and after (red) application of forskolin calculated from the fitting shown. **C**, Relative peak conductance (G_Na_^+^_max_, left), voltage for half-maximal activation (V_1/2_, middle), and slop factor (k, right) of Na^+^ channels, before and after (connected by lines) application of forskolin calculated from the recorded dataset in A. n = 7 axons. G_Na_^+^_max_, p = 0.00075; V_1/2_, p = 0.36; k, p =0.272

**Supplementary Figure 6.**
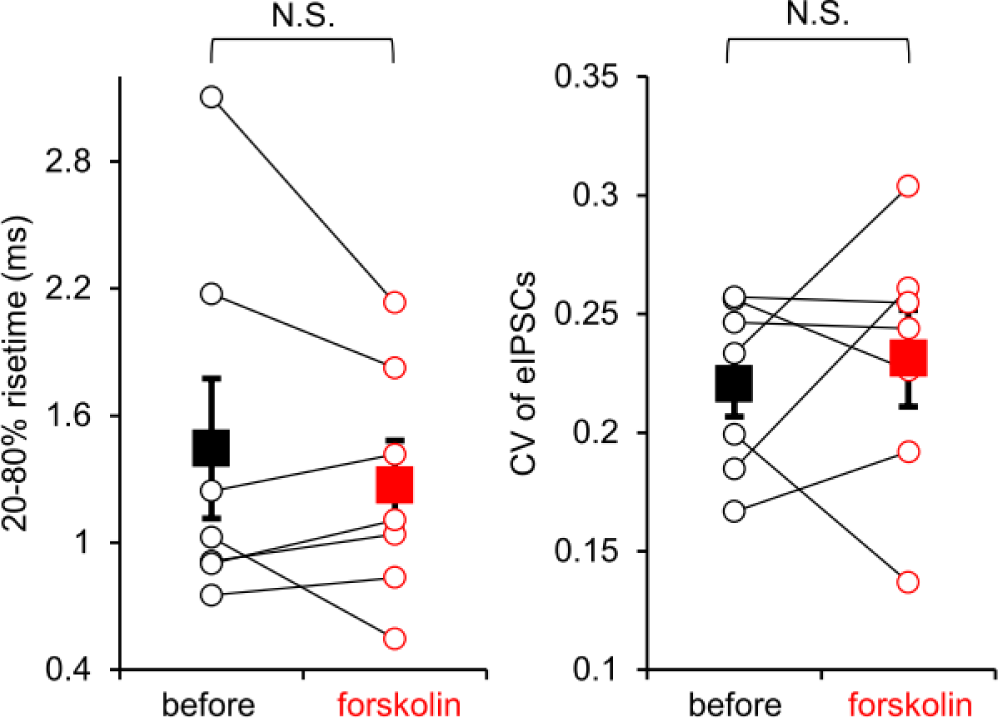
20-80% risetime (left) and CV (right) of eIPSCs of individual cells (open circles) and mean ± SEM (closed squares) before and after (connected by lines) application of forskolin. n = 7. risetime, p = 0.34; CV, p=0.60

**Supplementary Table 1.** Summary for recordings of axonal voltage-gated currents without (control) or with cAMPS-Sp.

|  | control (n =6) | cAMPS-Sp (n =6) | p value |
| --- | --- | --- | --- |
| I<sub>Na</sub><sup>+</sup> peak amplitude | 1.76 ± 0.27 nA | 0.99 ± 0.16 nA | *0.033 |
| I<sub>K</sub><sup>+</sup> peak amplitude | 4.23 ± 0.46 nA | 3.42 ± 0.82 nA | 0.409 |
| leak current | -31.2 ± 6.5 pA | -95.2 ± 14.3 pA | **0.0023 |
| apparent membrane resistance | 362.0 ± 55.7 MΩ | 149.1 ± 28.1 MΩ | **0.0066 |
| series resistance | 58.6 ± 2.3 MΩ | 69.2 ± 4.6 MΩ | 0.067 |
| soma-axon distance | 358.8 ± 78.7 μm | 377.5 ± 54.5 μm | 0.849 |

## Notes

### Competing Interest Statement

The authors have declared no competing interest.

